# Targeted Viromes and Total Metagenomes Capture Distinct Components of Bee Gut Phage Communities

**DOI:** 10.1101/2024.02.12.579852

**Authors:** Dino Lorenzo Sbardellati, Rachel Lee Vannette

## Abstract

Despite being among the most abundant biological entities on earth, bacteriophage (phage) remain an understudied component of host-associated systems. One limitation to studying host-associated phage is the lack of consensus on methods for sampling phage communities. Here, we compare paired total metagenomes and viral size fraction metagenomes (viromes) as methods for investigating the dsDNA viral communities associated with the GI tract of two bee species: the European honey bee *Apis mellifera* and the eastern bumble bee *Bombus impatiens*. We find that viromes successfully enriched for phage, thereby increasing phage recovery, but only in honey bees. In contrast, for bumble bees, total metagenomes recovered greater phage diversity. Across both bee species, viromes better sampled low abundance and low occupancy phage, while total metagenomes were biased towards sampling temperate phage and the most prominent phage. Additionally, many of the phage captured by total metagenomes were absent altogether from viromes. Comparing between bees, we show that phage communities in commercially reared bumble bees are significantly reduced in diversity compared to honey bees, likely reflecting differences in bacterial titer and diversity. In a broader context, these results highlight the complementary nature of total metagenomes and targeted viromes, especially when applied to host-associated environments. Overall, we suggest that studies interested in assessing total communities of host-associated phage should consider using both approaches. However, given the constraints of virome sampling, total metagenomes may serve to sample phage communities with the understanding that they will preferentially sample dominant and temperate phage.

## Introduction

Bacteriophage (phage) are the viruses which target bacteria. Phage are hypothesized to be among the most abundant biological entities on earth^1–3^ and to be important modulators of microbial communities ^2,4^. In marine and soil environments, where these prokaryotic viruses are relatively well studied, phage shape the microbiomes they infect by lysing dominant bacteria ^5–7^, influencing nutrient turnover and cycling^8,9^, and vectoring functional genes between bacterial hosts^10–13^. While the basis for much of our understanding of phage ecology stems from the study of marine and soil environments, there is a growing interest in understanding the role of phage in host-associated systems^14,15^. However, host-associated environments and free-living soil and marine environments represent different microbial ecosystems with different selective pressures. As such, insights gained from studying free-living phage communities may not always apply to communities of host-associated phage. Specifically, while the best methods for sampling free-living phage communities have been empirically compared^16,17^, little work has evaluated the best methods for sampling host-associated phage^18^. Moreover, given the importance of microbiomes to host health, understanding phage community dynamics and how phage differ between related animal hosts is a key priority.

Like their environmental counterparts, phage infecting host-associated bacteria (hereafter ‘host-associated’ phage) can affect the abundance, composition, and function of host-derived microbial communities. However, unlike free-living phage, host-associated phage must also adapt to their close association with a host and its microbiome. For example, the comparatively high-density bacterial communities associated with the human gut^19,20^, mucosal layer of corals^21^, and other host environments is suggested to favor integrase encoding temperate phage, as opposed to obligately lytic phage^22–24^. Phage targeting bacteria adhered to gut mucosa have also been shown to evolve the ability to adhere to and persist in the animal mucosa where their target bacteria reside^25^. Given the fundamental differences in the ecology of free-living and host-associated phage communities^22^, it remains unclear whether sampling methods developed for free-living communities are also appropriate for other habitat types.

Two methods are frequently used for describing dsDNA viral communities: bioinformatic mining of total metagenomes and targeted viral enrichments (viromes). In bioinformatic mining of total metagenomes, total genomic DNA is extracted from a sample, amplified, sequenced using ‘shotgun’ or untargeted methods, assembled into metagenomes, and phage computationally mined^26,27^. This approach is cheap, easy to perform, and offers simultaneous characterization of all DNA in a sample (including bacteria). However, only a small minority of the total metagenomic data generated will originate from phage, resulting in a relatively shallow sampling of phage communities. An alternative method, targeted virome sequencing, leverages the physical characteristics of viruses to select for the phage and virus-like particle fraction of a sample prior to nucleic acid extraction and sequencing^28,29^. With more sequencing devoted to viruses, targeted viromes can recover a greater diversity of phage relative to total metagenomes. This is illustrated by previous comparisons of viromes and total metagenomes in soil and freshwater samples^16,17^. However, whether viromes always outperform total metagenomes is not clear. Work applying these two methods to host-associated (human gut) and low-biomass (deep-sea marine) samples has produced conflicting results^17,18^. One explanation for these discrepancies is the biases associated with total metagenomes and targeted viromes^30,31^. For example, viromes remove bacterial cells prior to sequencing and, as a result, miss integrated temperate phage^32,33^. This is especially relevant in host-associated systems where temperate phage are abundant^32^. Total metagenomes may also be preferable to viromes logistically, given that viromes demand a relatively large biomass and are labor-intensive to generate, which can limit sample sizes in low-biomass systems.

Insects have become valuable models for exploring host-microbe interactions^34–36^. Social bees in particular house a simple (5 - 9 taxa), highly conserved, socially transmitted, gut bacterial community^37,38^ which contributes to host nutrient acquisition and pathogen defense^39–41^. These features position social bees as an excellent model for studying not only host-microbe interactions^42,43^, but tripartite host-microbe-phage interactions as well^44^. A series of recent studies using targeted virome approaches have shown that honey bees host a diversity of novel phage which target core bee gut bacteria^45–47^. While this work has advanced our understanding of bee phage, no work has evaluated how sampling method influences recovered bee phage communities. Given the large biomass required for viromes, total metagenomes may be advantageous for surveying the phage communities associated with bees and other biomass limited systems. Additionally, describing the phage associated with other social bees will develop our understanding of how phage communities differ between related hosts. Lastly, broader work comparing the role and diversity of phage among small invertebrate hosts will hinge on appropriate sampling methods, so comparing the performance of viromes and total metagenomics is particularly important for enabling future comparative study.

Here, we evaluate the phage communities inferred by applying total metagenomes and targeted viromes to managed *Apis mellifera* and commercially raised *Bombus impatiens* gut material. We hypothesize that targeted viromes will enrich for phage sequences and capture a greater diversity of phage, relative to bioinformatic mining of total metagenomes, and that phage will differ between host bee species. To interrogate these hypotheses, we first examine how host bee species and sampling method impact sequencing read data and viral enrichment. We then validate our sampling and computational methodology through rarefaction and by comparing our phage sequences to those previously described in honey bees. Next, to test how phage community differs across host bee and sampling methods, we compare phage community diversity, structure, and composition. We then use gene content, occupancy-abundance plots, qPCR, and bacterial community profiling to delve into what contributes to the apparent differences in phage communities. Overall, we find that honey bees host more phage than do bumble bees, but that the method which captured the most phage differed by bee species. This suggests that viromes and total metagenomes are each biased in their own way and appear to sample different populations of phage. As a result, we propose these techniques are complementary in describing the full diversity of host-associated phage.

## Methods

### Bee rearing, sampling, and experimental design

A detailed description of our methods (including sample collection, and bench and computational work) is provided in the supplemental materials. Briefly, bees were harvested from three colonies of managed honey bees (*Apis mellifera*) and three colonies of commercially reared bumble bees (*Bombus impatiens*, Koppert Biological Systems; Howell, MI, USA). Bumble bee colonies were maintained in the laboratory and provided with honey bee collected pollen and artificial nectar (Koppert Biological Systems) *ad libitum*, as per manufacturer’s recommendation. Honey bees were sampled from colonies maintained at the University of California, Davis apiary and were collected from inside the colony using a handheld bee vacuum between January 30^th^ – February 13^th^ 2023, 8:00am – 10:00am.

Bees were anesthetized via a 60 sec CO_2_ exposure and then euthanized via decapitation. Immediately following sacrifice, the mid-hindgut section was dissected from 100 bees (all collected from the same colony) and pooled for same day virome extraction. Bees destined for total metagenomic DNA extraction were collected the same day as virome samples and were stored at -20°C pending mid-hindgut dissection on the following day. All dissections took place in sterile PBS using ethanol and flame sterilized forceps. Each honey and bumble bee colony generated one targeted virome and three total metagenomes (**Fig. 1**).

**Figure 1:**
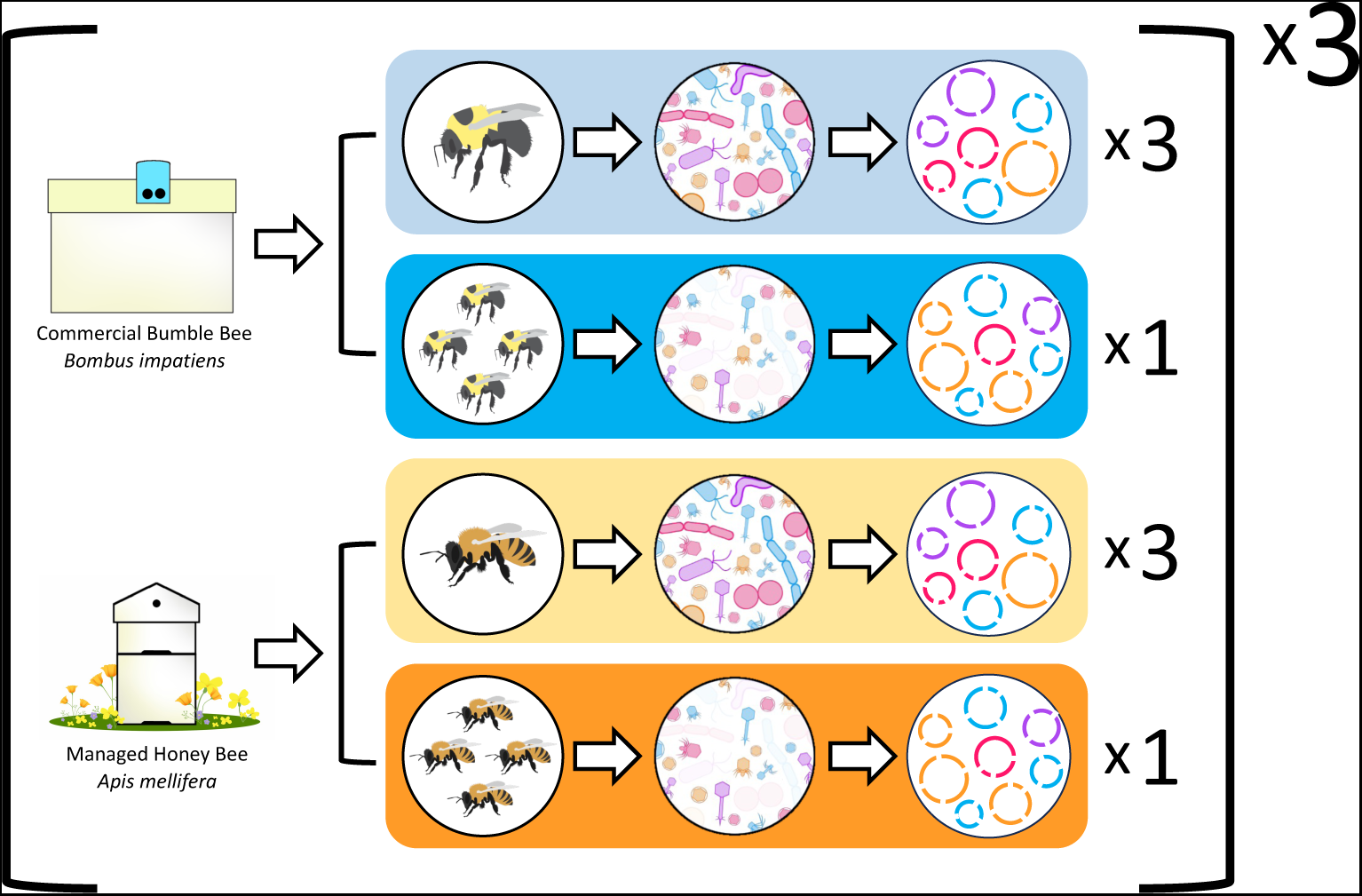
Graphical representation of the sampling scheme and methods used in this research. A total of three bumble bee and three honey bee colonies were sampled. From each colony of bee, we generated three total metagenomes and one targeted virome. Total metagenomes were sampled from individual bees, while targeted viromes were produced from the pooled guts of 100 bees. This sampling resulted in nine bumble bee total metagenomes (light blue), three bumble bee viromes (dark blue), nine honey bee total metagenomes (light orange), and three honey bee viromes (dark orange).

### Sample preparation and DNA extraction

Phage enrichments for targeted virome sequencing were carried out following a protocol adapted from previous publications^28,29,48^ and is described fully in the supplement methods. Briefly, bee gut pools were combined with 10 ml of Protein-supplemented phosphate-buffered saline (PPBS: 2% BSA, 10% PBS, 1% K-citrate, 150 mM MgSO4), homogenized, and submitted to a series of washes aimed at eluting phage particles. Phage were enriched via centrifugation, 0.22 um filtration, and ultracentrifugation. Lastly, phage pellets were resuspended in 200 uL water and DNase treated.

For total metagenomes, mid-hind guts of individual bees were dissected and homogenized and used for extraction. DNA was extracted from both virome and total metagenome samples using a DNeasy PowerSoil Pro kit (Qiagen, Hilden, Germany) following the manufacturer’s instructions with the optional Qiagen Vortex Adapter 15 min bead beating step.

### Library construction and DNA sequencing

Extracted DNA from samples and kit blank negative controls was submitted to the University of California Davis Genome Center for library prep and paired end 150 bp shotgun metagenomic sequencing. Libraries were prepared using a Kapa Hyper prep kit (Kapa Biosystems-Roche, Basel, Switzerland) with Illumina TruSeq adapters (Illumina, San Diego, CA) and a target insert size of 350 bp. Sequencing took place on an Illumina NovaSeq. All samples were sequenced on the same run.

### Read processing, viral prediction, and vOTU table construction

Briefly, all demultiplexed *.fastq.gz* files from the Davis Genome Center were trimmed and quality filtered using Trim-Galore^49^ and Trimmomatic^50^. Reads aligning to negative samples or the genomes of *A. mellifera* and *B. impatiens* were removed using Bowtie2^51^. Cleaned reads were k-merized for complexity comparisons using Sourmash^52^. The program metaSPADES was used to construct assemblies. Assemblies ≥5 kb were passed to Virsorter2^53^ for phage prediction. Phage with ≥95% nucleotide identity were collapsed into viral operational taxonomic units (vOTUs) using CD-HIT^54^ and were annotated with Prokka^55^ using the PHROGs database^56^. vOTUs were classified as temperate when they were predicted to encode an integrase gene. Finally, CoverM^57^ was used to map cleaned reads against vOTUs. The resulting coverage table was normalized to coverage per million reads.

### Bacterial community and density measurements

Bacterial communities were predicted directly from cleaned total metagenomic reads with Kraken2^58^ and Bracken^59^ using default parameters. Bacterial copy number was quantified from the DNA extracts used for metagenome sequencing via a qPCR protocol adapted from Christensen et al^60^. Each master mix solution contained: 5ul SSO Advanced Universal SYBR Supermix 41 (Biorad, Hercules, CA, USA), 0.3ul of primers (10uM) targeting the 16S region of the rRNA gene (799F= 5’-AACMGGATTAGATACCCKG-3’;1115R= 37 5’-AGGGTTGCGCTCGTTG-3’), 3.4ul Molecular grade water, and 1ul of template DNA (diluted 1:1000 in molecular grade water). Reactions were performed in triplicate for each sample.

### Statistical and ecological analyses

Statistical analyses were conducted in R^61^ v4.2.3. Briefly, alpha diversity, beta diversity, and PERMANOVAs were compared between host and sampling method using vegan^62^ and phyloseq^63^. Linear mixed effect models were built using lme4^64^, post-hoc tests were run using emmeans^65^, and T-tests were performed using base R. Gene-sharing networks were generated outside of R using the program vConTACT2^66,67^. Genome alignment plots were constructed using Clinker^68^.

Phage-host predictions were performed using a custom CRISPR-spacer analysis. In short, we built bacterial metagenome assembled genomes (MAGs) from our total metagenomic dataset using metaSPADES and MetaBat2^69^, combined these MAGs with NCBI assemblies of common bee gut bacteria, mined all these sequences for CRISPRs using MINCED^70^, and then aligned those CRISPRs to our vOTUs using Blastn^71^. Lastly, to improve the number of vOTUs with host assignments, we combined phage-host assignments generated by our CRISPR analysis with the clusters predicted by our vConTACT2 network to infer the hosts of whole phage clusters, following Rosso et al^45^. Bacterial copy number was compared between species using linear models.

## Results

### Bumble bee viromes produce fewer high-quality reads than honey bee viromes

Our sequencing effort generated a total of 419,884,845 raw read pairs, 2.47% (10,385,967) of which came from extraction negatives. While the number of raw read pairs did not differ significantly between samples (**Fig. S1;** Bee t_5.40_=0.54 p=0.61, Type t_8.47e23_=1.58 p=0.11, Bee:Type t_1.07e23_=-1.38 p>0.17), there were differences in the number of high-quality read pairs remaining after quality control (Bee t_5.81_=2.87 p<0.05, Type t_2.75e25_=-0.33 p=0.74, Bee:Type t_2.75e25_=4.69 p<0.001) (**Fig. 2A**). After filtering, honey bee viromes averaged 15.56 million high-quality read pairs per sample, while bumble bee viromes averaged only 5.49. Total metagenome samples generated more similar numbers of high-quality reads per sample, with an average of 8.59 and 5.85 million read pairs generated from honey and bumble bee samples, respectively.

**Figure 2:**
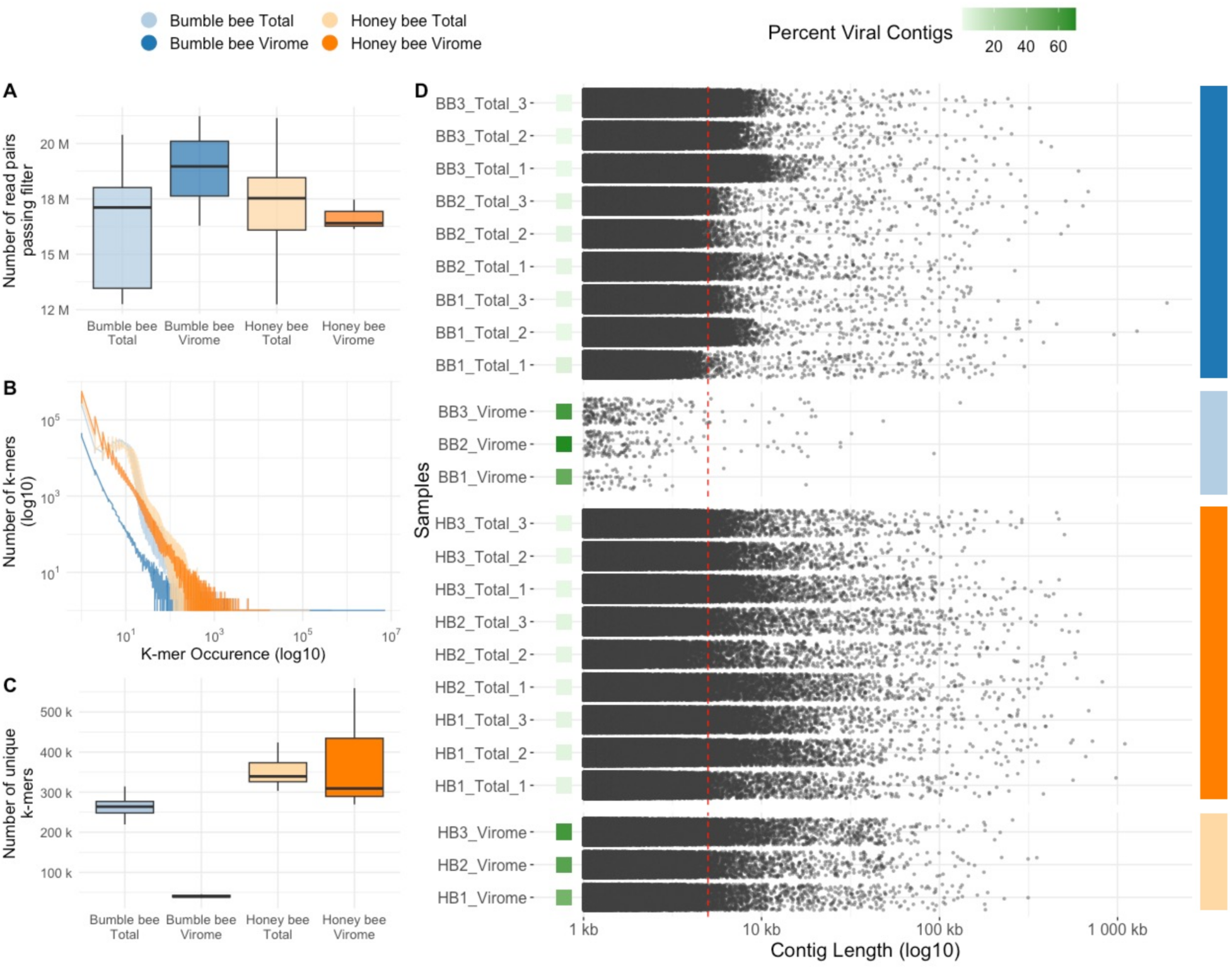
Figures describing sequencing and assembly quality. Color denotes sample groups. **A)** Boxplots describing sequencing depth. The x-axis shows sample groups. The y-axis displays the total number of high-quality reads produced. **B)** Line plots describing sequence complexity of the reads shown in panel **A**, as measured by the frequency distributions of 31bp sized k-mers. The x-axis presents k-mer occurrence (i.e. how abundant a particular k-mer was in a given sample’s read), while the y-axis shows the number of k-mers with a certain occurrence. **C)** Boxplots describing the number of unique 31bp sized k-mers present in the read libraries of each group of samples. The x-axis shows sample groups. The y-axis shows the number of unique k-mers. **D)** Jittered dot-plots describing the length distributions of contigs assembled from each sample. Only contigs >= 1Kbp bp are shown. Individual samples are shown on the y-axis. The x-axis shows contig length. Each point represents a single contig. A dotted red line is drawn at 5Kbp. For each sample, a green square describes phage enrichment (the number of phage identified, divided by number of contigs assessed for phage, times 100).

### Bumble bee virome reads are less complex than other samples

Next, we measured the complexity of read libraries to further investigate how bee species and sampling method influenced the data we generated. This was done by comparing the number and occurrence of 31bp sized k-mers amongst each set of high-quality read libraries (**Fig. 2B**). The results of this analysis suggest that, across the range of k-mer occurrences, bumble bee viromes consistently contain fewer k-mers of a given occurrence than did other sample types. Moreover, viewing only the y-intercept of this plot (**Fig. 2C**) shows that bumble bee viromes contain a lower number of singleton k-mers, relative to all other sample types (Bee t_5.34_=2.49 p=0.052, Type t_16_=-6.47 p<0.001, t_16_=5.14 p<0.001,). This decreased number of singleton k-mers, and the tendency towards lower k-mer occurrences suggest that bumble bee viromes are less complex compared to the other sample types in this dataset.

### Targeted virome assemblies are enriched in phage

To evaluate the size and number of assemblies produced from each read library, and to quantify the extent to which viromes enriched for phage sequences, we visualized the distribution of assembly sizes in each sample, as well as the percentage of assemblies in a sample predicted to encode phage sequences (**Fig. 2D**). A total of 4,278 and 37 contigs ≥5kb were assembled from honey and bumble bee viromes, respectively. After passing to VirSorter2, these contigs yielded 2,382 honey bee and 24 bumble bee putative phage sequences. Meanwhile, honey and bumble bee total metagenomes produced 16,615 and 14,335 contigs ≥5kb, respectively, which yielded 404 and 140 putative phage sequences. This means that 53.94% of honey bee and 60.57% of bumble bee virome contigs ≥5kb were predicted to be viral, while only 2.42% and 2.13% of honey and bumble bee total metagenome contigs ≥5kb were annotated as viral. The enrichment of phage sequences in viromes, relative to total metagenomes, provide validation that viromes successfully enrich for phage when applied to bee guts.

Next, we used rarefaction to assess if the different number of putative phage sequences recovered by honey and bumble bee viromes (**Fig. 2D**) was a result of differences in sampling depth **(Fig. 2A**). This analysis shows that even at equal sampling depths, honey bee viromes consistently recover >40x the number of phage recovered from bumble bee viromes (**Fig. S2A and S2B**).

### Phage target core bee bacteria and resemble previously described honey bee phage

After measuring the ability of viromes to enrich for phage sequences, we sought to characterize the phage communities we detected in terms of similarity to previously described bee phage, putative bacterial hosts, and predicted taxonomy. To do this, we first constructed vOTUs by collapsing putative phage sequences at a 95% average nucleotide identity. This reduced the number of putative phage from 2,950 to 2,560. We then built gene-sharing networks comparing our vOTUs to those previously described by other bee phage studies (**Fig. 3A**). In this network, many of the vOTUs from the current study clustered with those described by other bee phage studies, suggesting they are related at roughly the genus level^66,67^. For example, viral cluster 29 contains sequences recovered by the current study, Rosso et al^45^, and Deboutte et al^46^ (**Fig. S3**). We interpret these results as further validating our computational methodologies and further supporting the result that bumble bees host reduced diversity phage communities.

**Figure 3:**
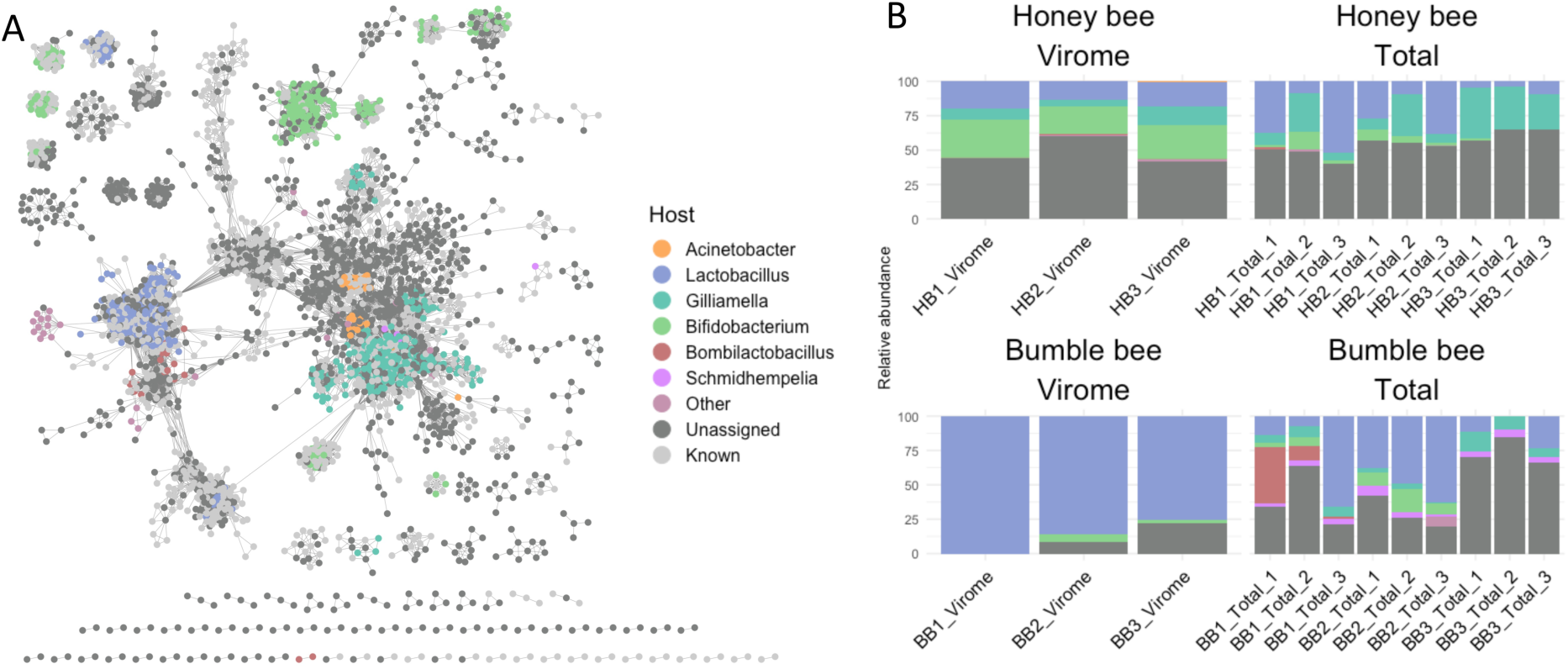
Figures describing phage identified in our dataset, relative to those identified previously, and the overall phage communities found in individual samples. **A)** Weighted gene-sharing network of all non-singleton vOTUs identified in our study. Individual nodes are phage. Nodes are connected by edges when phage share genes. Nodes are colored based on the predicted bacterial host of phage. Phage with “Known” as a predicted host are previously described bee phage. **B)** Stacked bar-plot describing the community of phage found in each sample based on predicted host. Both panel **A** and **B** use the same color palette.

Next, we assigned bacterial hosts to our vOTUs. To focus this and all downstream analyses on only the most abundant phage, we filtered vOTUs to retain only those with a coverage ≥5, resulting in 2,481 vOTUs across 24 samples. Overall, we assigned hosts to 29.91% (742) of the 2,481 vOTUs appearing in our dataset (**Fig. 3B**). In terms of relative abundance, phage with assigned hosts made up an average of 89.34% and 44.75% of bumble bee viromes and total metagenomes and 50.54% and 43.89% of honey bee viromes and total metagenomes, respectively. Notably, the high proportion of phage with assigned hosts in bumble bee viromes is likely confounded by the fact that these samples had an extraordinarily diminished phage diversity.

Phage predicted to target the core social bee bacterial genera *Lactobacillus*, *Bifidobacterium*, and *Gilliamella* were the most abundant of those with identified hosts in our dataset (**Fig. 3B**). However, the relative abundance of phage targeting these genera varied by bee species and sampling method (**Table S1)**. Phage targeting *Lactobacillus* were most abundant in bumble bees, especially bumble bee viromes, while *Bifidobacterium* targeting phage were most abundant in honey bee samples, particularly viromes. *Gilliamella* phage were not recovered in bumble bee viromes, but were found in most bumble bee total metagenomes, and were approximately 2x more abundant in honey bee total metagenomes than they were in honey bee viromes. Phage targeting the core bumble bee symbiont *Schmidhempelia* were exclusively found in bumble bee total metagenomes. We also found signatures of colony-level variation, for example phage targeting *Bombilactobacillus* were detected in metagenomes from individuals sampled from colony 1, but no individuals from colonies 2 or 3.

We assigned family level taxonomy to our phage if they clustered with reference phage in gene-sharing networks. In total, we only assigned taxonomy to 9.83% (244) of 2,481 vOTUs (**Fig. S4**). The large majority of classified vOTUs belonged to Siphoviridae (104), Myoviridae (94), and Podoviridae (40), with a smaller number being classified as Autographiviridae (3) and Zobellviridae (1) and multiple assignment (2).

### Bee species differ in phage diversity and composition

Next we tested if vOTU diversity and community composition differed between bee species and sampling method. While the diversity and richness of communities followed similar trends (**Fig. 4A and 4B**), there were substantial quantitative differences between these two metrics. When we compared Shannon’s diversity, bee species, sampling method, and their interaction all significantly affected vOTU diversity (**Fig. 4A**; Bee t_20_=2.315, p=0.0313; Type t_20_=-5.992, p<0.001; Bee:Type t_20_=8.004, p<0.001). Meanwhile, only the interaction between bee species and sampling method significantly impacted community richness (**Fig. 4B**; Bee t_20_=0.690, p=0.498; Type t_20_=-0.160, p=0.874; Bee:Type t_20_=10.227, p<0.001). Overall, these results suggest that sampling method can significantly influence the alpha diversity of inferred phage communities, but that this can differ in magnitude and direction according to bee species.

**Figure 4:**
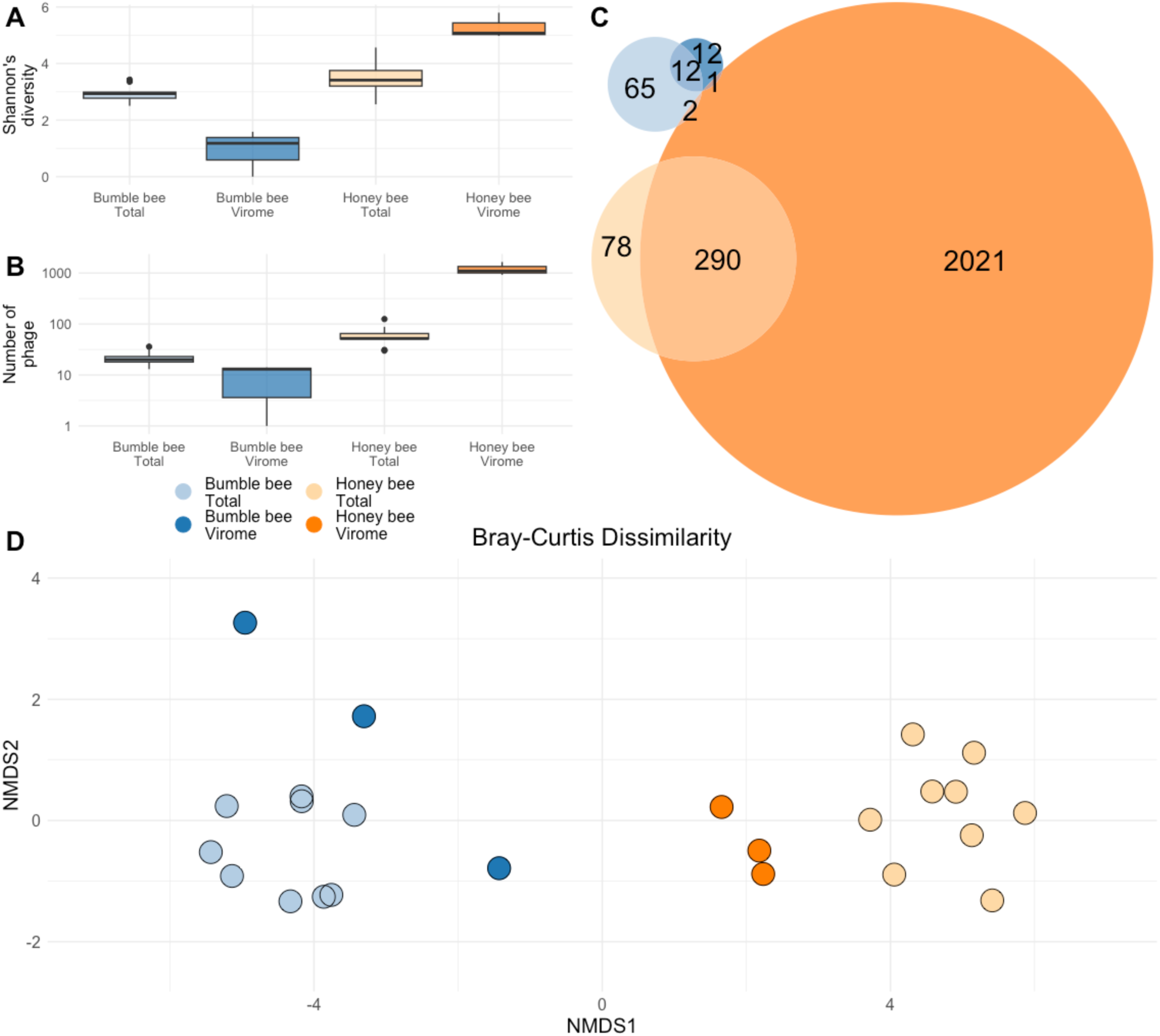
Figures describing the alpha and beta diversity of the phage communities identified in our samples. Blue represent samples taken from bumble bees. Orange represents samples taken from honey bees. Lighter colors are total metagenomes. Darker colors are viromes. **A)** Boxplots describing the **A)** Shannon’s diversity and **B)** vOTU richness associated with each of our sample types. The x-axis groups samples. The y-axis shows diversity and richness scores. **C)** Euler diagram describing phage community overlaps between each of our sample types. Numbers correspond to the number of phage present in each section of the graph (i.e. 290 phage were found in both honey bee viromes and total metagenomes). Circle size is proportional to number of phage. **C)** Non-metric multidimensional scaling (NMDS) ordination describing the Bray-Curtis dissimilarity of all samples on our dataset. Individual points are samples. Color represents sample type. The closer two points are to each other, the more similar they are in phage community.

In honey bees, 12.1% (290/2,392) of all vOTUs were shared between virome and total metagenome samples. Similarly, in bumble bees, 12.9% (12/93) of vOTUs were detected by both sampling methods. Meanwhile, only a few vOTUs were detected in both bee species (0.121%; **Fig. 4C**). Additionally, vOTU community composition was predicted primarily by host bee species, and secondarily based on sample method (**Figure 4D**; Bray-Curtis PERMANOVA Bee R^2^=0.143 F_1,20_=4.17 p<0.001; Method R^2^=0.0859 F_1,20_=2.506 p<0.001; Bee:Method R^2^=0.0859 F_1,20_=2.505 p<0.001).

### Bacterial community variation predicts phage community variation

We hypothesized that differences in phage richness and community composition between bee species may be driven by differences in bacterial abundance and composition. First, we used qPCR to compare the number of bacteria found in the guts of honey and bumble bees (**Fig. 5A**) and found that the average mid-hindgut 16S gene copy number was significantly lower in bumble bees than it was in honey bees (t_9.13_=-3.54 p<0.01).

**Figure 5:**
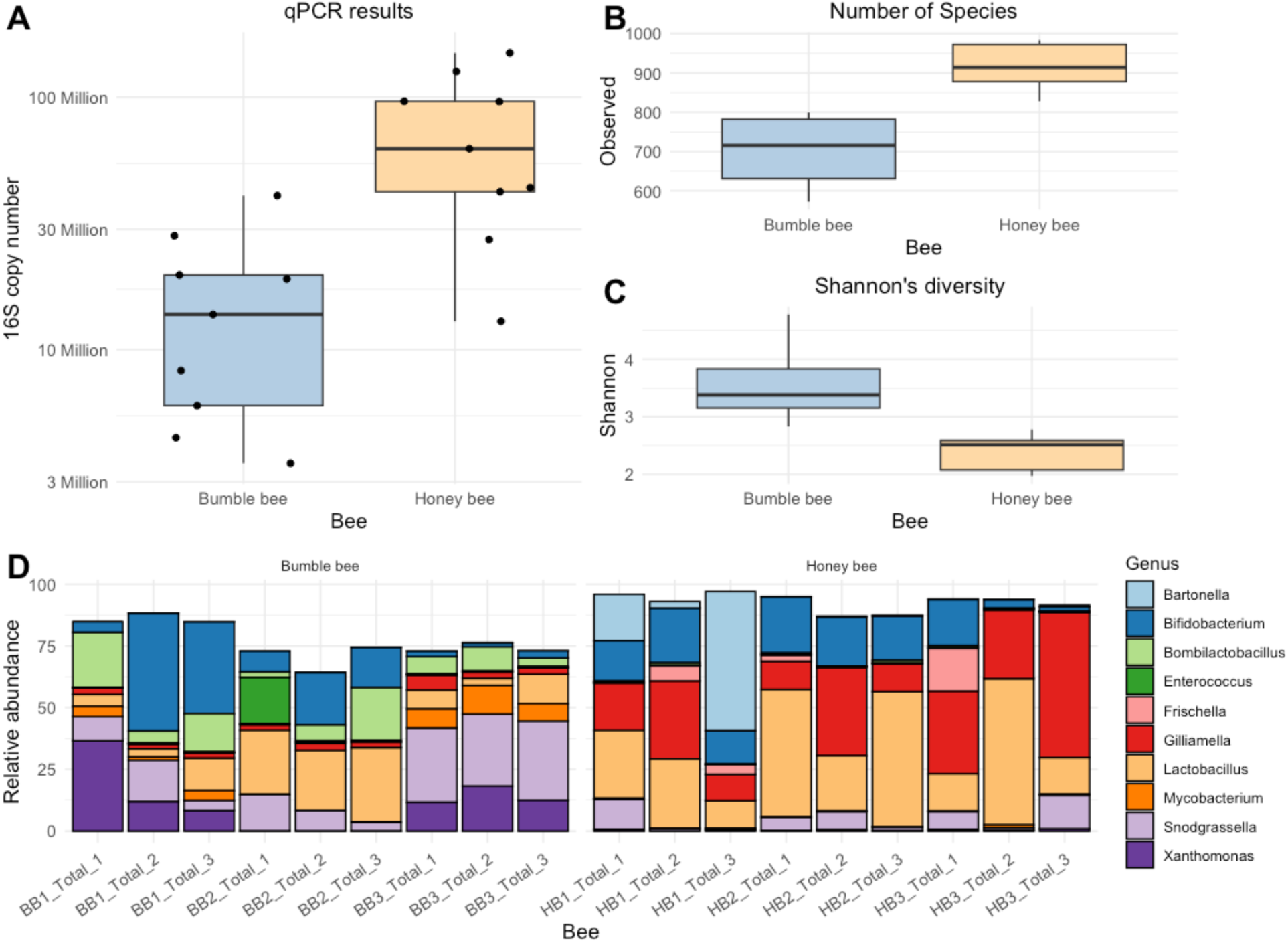
Honey and bumble bees differ in bacterial community density, diversity, and composition. **A)** Boxplots displaying the 16S rRNA gene copy number present in the mid-hindgut region of sampled honey and bumble bees. **B)** Boxplot presenting the number of observed bacterial species in honey and bumble bee. **C)** Boxplot showing the species level Shannon’s diversity of honey and bumble bee gut bacterial communities. **D)** Stacked bar plot describing honey and bumble bee bacterial communities at the genus level.

We then used Kraken2 and Bracken to estimate the diversity and taxonomic profile of bacterial communities (**Fig. 5B, 5C, 5D**). At the species level, bumble bee bacterial communities hosted lower bacterial richness and greater Shannon’s diversity than honey bee bacterial communities (Richness t_13.78_=-6.27 p<0.001, Shannon t_11.70_=-6.27 p<0.001), suggesting fewer bacterial taxa were present in the bumble bees we sampled, but that bumble bee bacterial communities were more evenly divided amongst their constituent bacteria.

Lastly, we used a Mantel test to compare the relationship between bacterial and phage community structure (**Fig. S5**). We found a high degree of similarity between the bacterial and phage communities (R=0.74 p<0.001), meaning that individual bees with similar bacterial communities also have similar phage communities.

### Total metagenomes capture prophage but miss rare phage

After identifying factors which explain some of the host-specific differences in phage communities, we sought to evaluate why sampling method (virome vs total metagenome) also led to significant differences in inferred phage communities. For this, we first compared the gene content of vOTUs recovered from viromes and total metagenomes. Because total metagenomes are primarily comprised of bacterial DNA, we hypothesized that this sampling method would enrich for temperate phage. Supporting this hypothesis, we found that 189 of the 2075 (9.11%) vOTUs identified in viromes were predicted to encode an integrase gene, while 89 of the 406 (21.92%) vOTUs recovered from total metagenomes encoded an integrase (**Fig. S6**). Moreover, integrase encoding phage had a higher relative abundance in total metagenomes than in viromes (**Fig. 6A**; Bee t_14.813_=-0.625 p=0.541; Type t_16_=2.845 p<0.05; Bee:Type Type t_16_=0.463 p=0.650). These results suggest that, relative to viromes, total metagenomes are biased towards sampling prophage.

**Figure 6:**
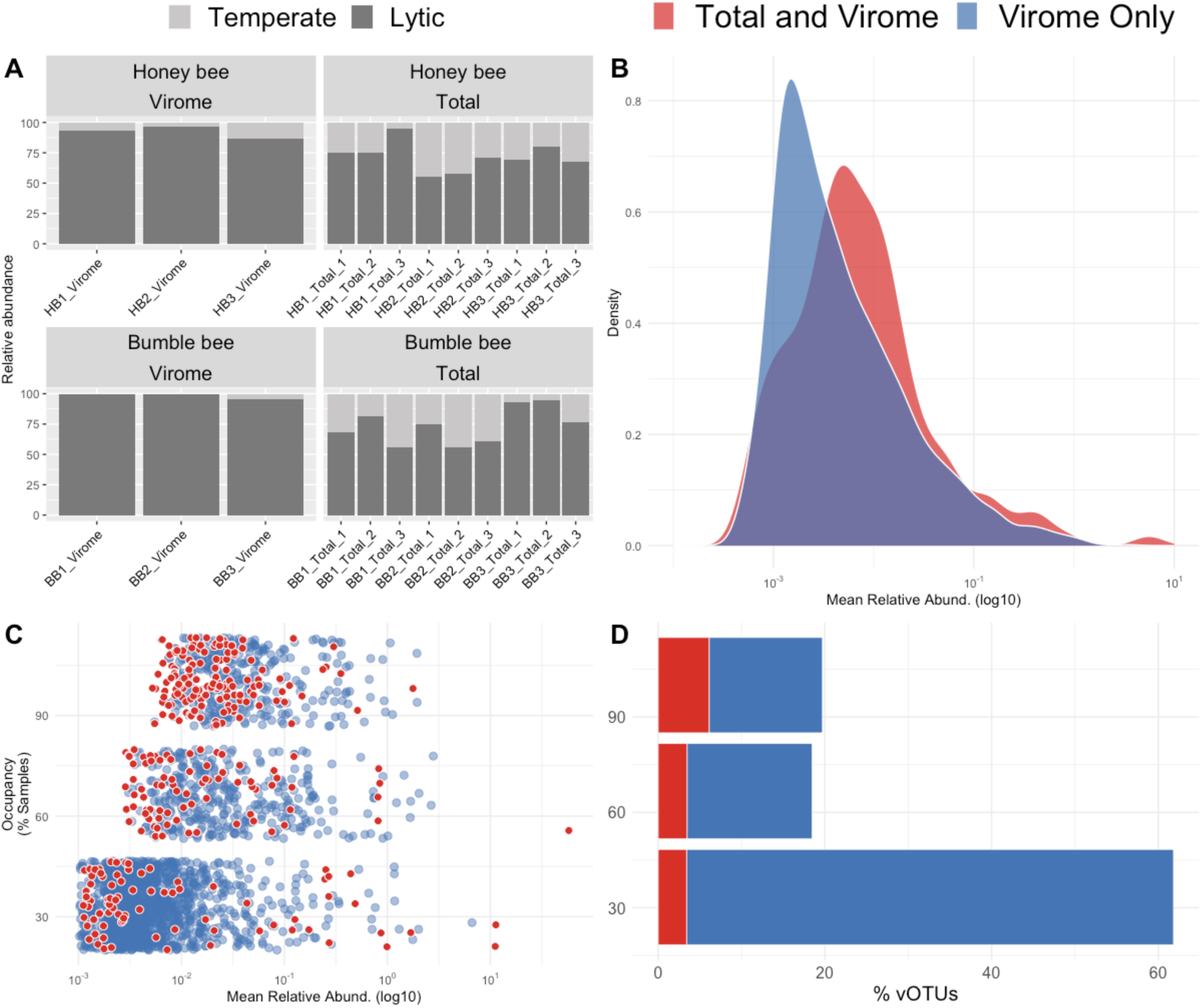
vOTUS sampled by viromes and total metagenomes differ in integrase content, occupancy, and abundance. **A)** Stacked bar plot showing the relative abundance of temperate and lytic vOTUs in virome and total metagenome samples. **B)** Density distributions showing the average relative abundance of all vOTUs found in viromes. vOTUs unique to viromes are shown in blue. vOTUs found in both viromes and total metagenomes are shown in red. **C)** Dot plot showing the occupancy-abundance relationship of all virome vOTUs. The x-axis describes average relative abundance, while the y-axis shows the percent occupancy of each vOTU. As before, vOTUs unique to viromes are shown in blue and those found in both viromes and total metagenomes are shown in red. **D)** Stacked bar plot describing the percent of vOTUs found in one, two, or three samples. The x-axis represents the percent of vOTUs. The y-axis the occupancy. Similar to B and C, vOTUs unique to viromes are shown in blue and those found in both viromes and total metagenomes are shown in red

Lastly, to examine how sampling methods differ in recovery of rare vs abundant phage, we visualized the occupancy-abundance relationships of virome-specific vOTUs vs vOTUs recovered by both methods (**Fig. 6B and 6C**). This analysis revealed that vOTUs identified in total metagenomes were among the most abundant and prevalent vOTUs found in viromes, suggesting that total metagenomes recover more abundant and prevalent phage, while potentially missing rarer and lower occupancy phage.

## Discussion

Although phage are integral members of complex microbiomes, methods for describing phage communities are still being developed. Here we compare how two common forms of phage community sampling, bioinformatic mining of total metagenomes and targeted sequencing of viromes, influence the phage communities recovered from the guts of two commercially important bee species: *Apis mellifera* and *Bombus impatiens*. Similar to the results of previous studies^16–18^, we show that sampling method significantly affected alpha diversity, beta diversity, composition, and structure of sampled phage communities. In particular, we show that viromes outperform total metagenomes in terms of the number of phage recovered, but that this can depend on the specific environment being sampled. Overall, viromes were better at sampling rare and less prevalent phage when bacterial and phage diversity was high, while total metagenomes captured a greater diversity of phage in low biomass samples. We also show that, regardless of sample biomass, total metagenomes were enriched for temperate phage, compared to viromes, and were consistently able to recover phage not found in viromes. Given these results, we suggest that viromes and total metagenomes each have limitations, are complementary, and that choice of one method over the other likely depends on the environment being sampled.

### Honey bees host a core phage community

Viewing our results within the broader context of bee phage, our analyses show that many features of the phage communities identified here resemble those previously described in honey bees sampled from North America and Europe^45–47^. For example, *Siphoviridae*, *Myoviridae* and *Podoviridae* were among the most common viral families that we and these previous studies identified. Similarly, *Lactobacillus*, *Gilliamella*, *Bifidobacterium*, and other core bee gut bacteria were among the most predicted phage hosts in our and previous studies. Notably, while previous bee phage studies found a number of phage predicted to target *Bartonella*, none of our phage were predicted to target this bacterial genus. This is likely because no available *Bartonella apis* assemblies met the strict criteria for inclusion in our CRISPR spacer analysis.

Delving deeper, our gene-sharing network and clinker analyses show that many of our phage share not only large genomic regions with previously described honey bee phage, but that gene synteny in these geographically disparate viral genomes is also conserved. This conservation between honey bee phage communities is similar to what was observed by Busby et al^47^. Likewise, Rosso et al^45^ showed that the genomes of honey bee-associated phage sampled from Europe are able to recruit reads from Japanese honey bee total metagenomic datasets.

Altogether, these findings support the idea that, similar to the highly conserved nature of some bee gut bacterial communities^37,38,43,72^, some features of bee gut phage communities are also conserved between bees sampled across geographically disparate populations. Future work should focus on identifying what features of phage communities, whether individual genes, clusters of genes, or perhaps whole genomes, are conserved between different populations of bees.

### Host bacterial communities predict phage communities

When compared to free-foraging managed honey bees, the commercially produced bumble bees we sampled hosted significantly lower diversity phage communities. Although differences in species biology and colony size may play a role, we attribute this low phage diversity primarily to the low density and diversity bacterial communities these bumble bees hosted, suggesting that phage diversity and abundance may track bacterial diversity and abundance across hosts. However, whether such results are generalizable to wild bumble bees is unclear. In our study, bacterial 16S rRNA copy number in bumble bee guts ranged from 10^6^ to 10^7^, whereas previous work in the same species reported 10^8^ - 10^9^ copies of the 16S rRNA gene^37,73–75^. Given we used standard rearing methods and diet, we hypothesize that lower bacterial titer may be due to the age of bees we sampled. Previous work in bumble bees shows that bacterial density increases with worker age, saturating approximately four days after worker emergence^73^. Future work could examine how phage abundance and composition changes through worker development and if this reflects previous observations of bacterial succession with worker age^17^. More generally, these findings suggest that features of host bacterial communities (i.e. density, diversity, and structure) may be used to predict phage community features. This is further supported by our Mantel analysis showing a correlation between bacterial and phage community dissimilarity.

### Viromes and total metagenomes reveal complementary phage communities

There are several possible reasons as to why the phage communities inferred by viromes and total metagenomes differ. One explanation is the way we generated viromes and how this influenced the specific population of phage sampled. Research by Hoyles et al.^76^ has shown that passing human fecal samples through a 0.22 um filter reduces the number of phage recovered by nearly half. As such, size filtration may have led some phage which are present in our bee guts to have been excluded from viromes. This size exclusion may explain why we, and previous virome vs total metagenome studies^16–18^, consistently show that some phage are only found in total metagenomes. The particular types of phage which are being excluded likely further explain the total metagenome and virome differences we observed. We found that putative temperate phage were more abundant and prevalent in total metagenomes than they were in viromes. This agrees with existing literature which suggests that size filtration can specifically remove integrated lysogenic phage, large jumbo phage, and phage adhered to bacterial cells and particles^31,33,77,78^.

While it is tempting to expand these results to different environments by stating total metagenomes always sample temperate phage missed by viromes, work in other systems implies a more nuanced reality. In agricultural soils, Medellin et al.^16^ identified only three phages that were present in total metagenomes and absent from viromes. This suggests that the degree to which sample processing skews viromes differs by environment. As a result, we suggest that the relative utility of viromes and total metagenomes likely depends on the environment being sampled. Samples with a high abundance of temperate phage, such as host-associated systems and low-biomass environments, may benefit from total metagenomes. This is similar to the conclusions made by Kosmopoulos et al.^17^ which suggest that the choice of viromes vs total metagenomes should be environment specific.

While our results highlight the ability of total metagenomes to recover phage missed by viromes, they also showcase the capacity of viromes to sample a greater diversity of phage. Similar to previous comparisons of viromes and total metagenomes in human gut, soil, and freshwater environments^17,18,29^, our honey bee viromes recovered a substantially larger number of phage than did total metagenomes. Further, the occupancy-abundance relationships examined here and by Medellin et al.^16^ show that total metagenomes tend to be biased towards sampling the most abundant and prevalent phage, while viromes are more successful at sampling rare phage. Taken together, these results support previous work documenting that viromes produce a deeper sampling of phage communities compared to total metagenomes^31,79,80^.

### Conclusions

By comparing bioinformatic mining of total metagenomes and targeted viromes across two bee species, we found that sampling method significantly affected inferred phage communities, but that the directionality of these differences can depend on the host being sampled. In general, total metagenomes tended to be biased towards sampling both putatively temperate phage and phage which were highly abundant. In samples with a relatively high biomass (e.g., honey bees), viromes produced a greater diversity of phage and a better sampling of rare phage. In contrast, when applied to relatively low biomass samples (e.g., bumble bees), total metagenomes captured a greater diversity of phage than did viromes. Regardless of sample biomass, there were always phage unique to both viromes and total metagenomes. In conclusion, we suggest that these methods are complementary and recommend that both be used to capture the full diversity of phage present in a gut environment. However, given that virome sampling is not always feasible (i.e. in the case of field collected insects) total metagenomes may serve to sample phage communities with the caveat that they will preferentially sample dominant and temperate phage.

## Acknowledgements

We thank Dr. Elina L. Nino, Joseph Tauzer, and the UC Davis Harry H. Laidlaw Jr. Honey Bee Research Facility staff for help sampling managed honey bee colonies, Samantha R. Eck and members of the Vannette, Emerson, and Brown labs for comments on the manuscript, Anoushka Basu for help dissecting bees and the UC Davis Genome center for sequencing.

## Data Availability

The raw sequencing data generated in this study is available on NCBI under the Bioproject accession number PRJNA1072153. All scripts used to process this data is publicly available at https://github.com/dsbard/Bee_phage_2023.

