## Supplemental Figures for "Targeted Viromes and Total Metagenomes Capture Distinct Components of Bee Gut Phage Communities"

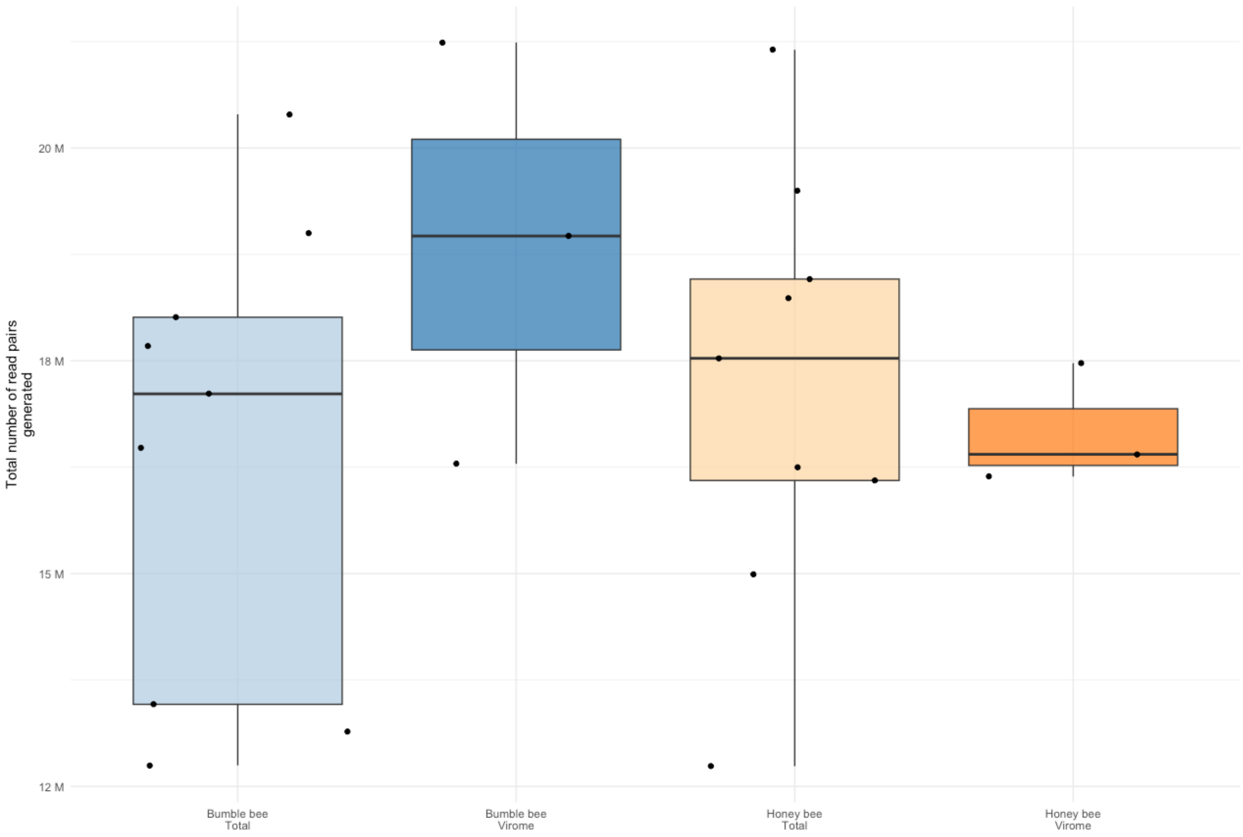


**Figure S1:** Boxplots describing the number of raw read pairs generated per sample type. Sample type is shown along the x-axis. Total number of reads pairs, in millions, generated is represented on the y-axis. All samples produce a roughly equal number of raw reads.

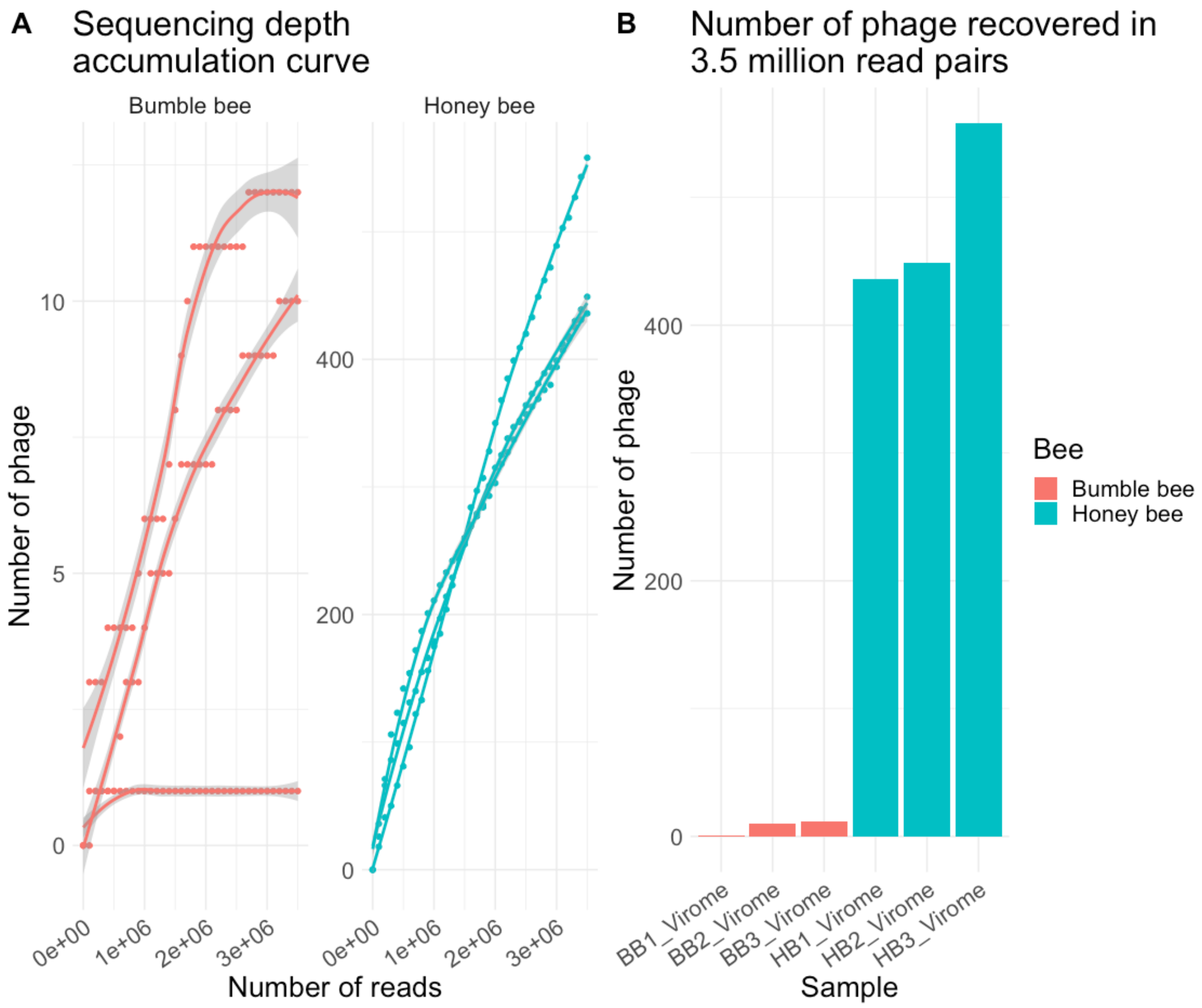


**Figure S2:** When sampled at equal depths, honey bees recover a greater number of phage sequences. **A)** Species accumulation curve showing the number of phage recovered from honey and bumble bees as the number of reads considered increases. **B)** The number of phage recovered from honey and bumble bees when sampled at an equal depth (3.5 million read pairs).

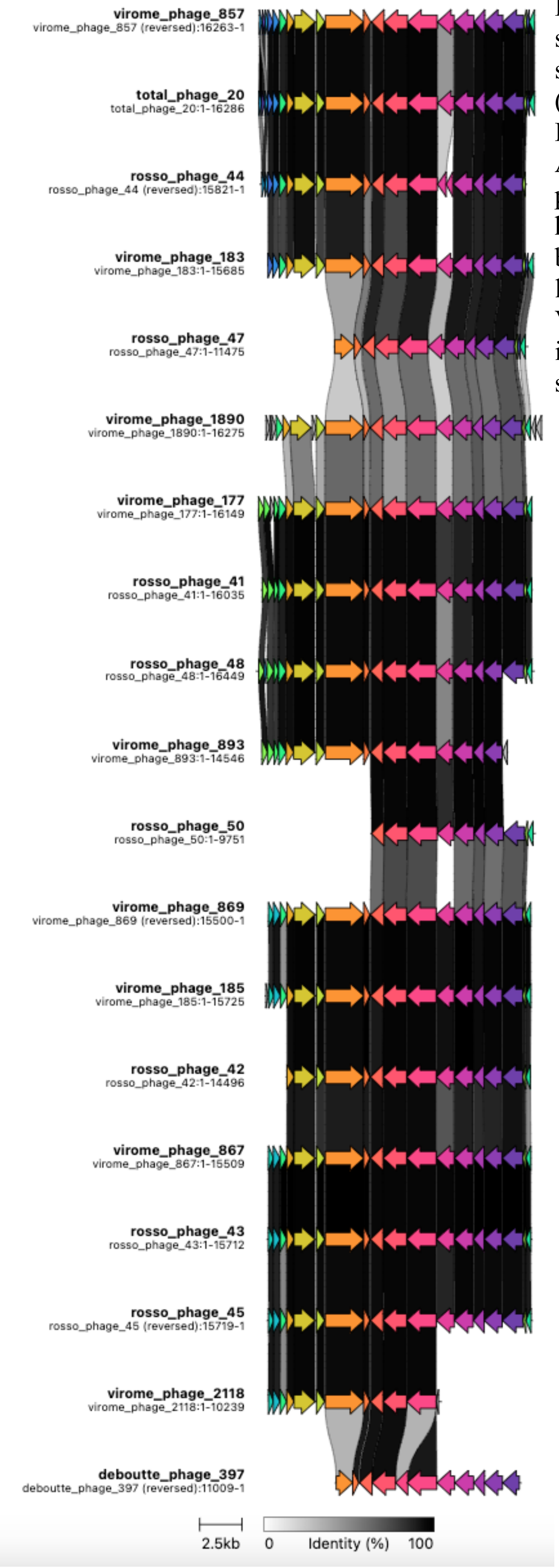


**Figure S3:** Clinker plot displaying gene sharing amongst all the phage in a single gene-sharing network cluster. The cluster shown (VC_29) is predicted to target *Lactobacillus*. Each row is an individual phage in this cluster. Arrows represent the genes present in those phage. Connections between phage represent homologous genes. Connections are colored based on average amino acid identity. Phage labeled with the study they were identified in. Virome and total metagenome phage were identified in our virome and total metagenome samples, respectively.

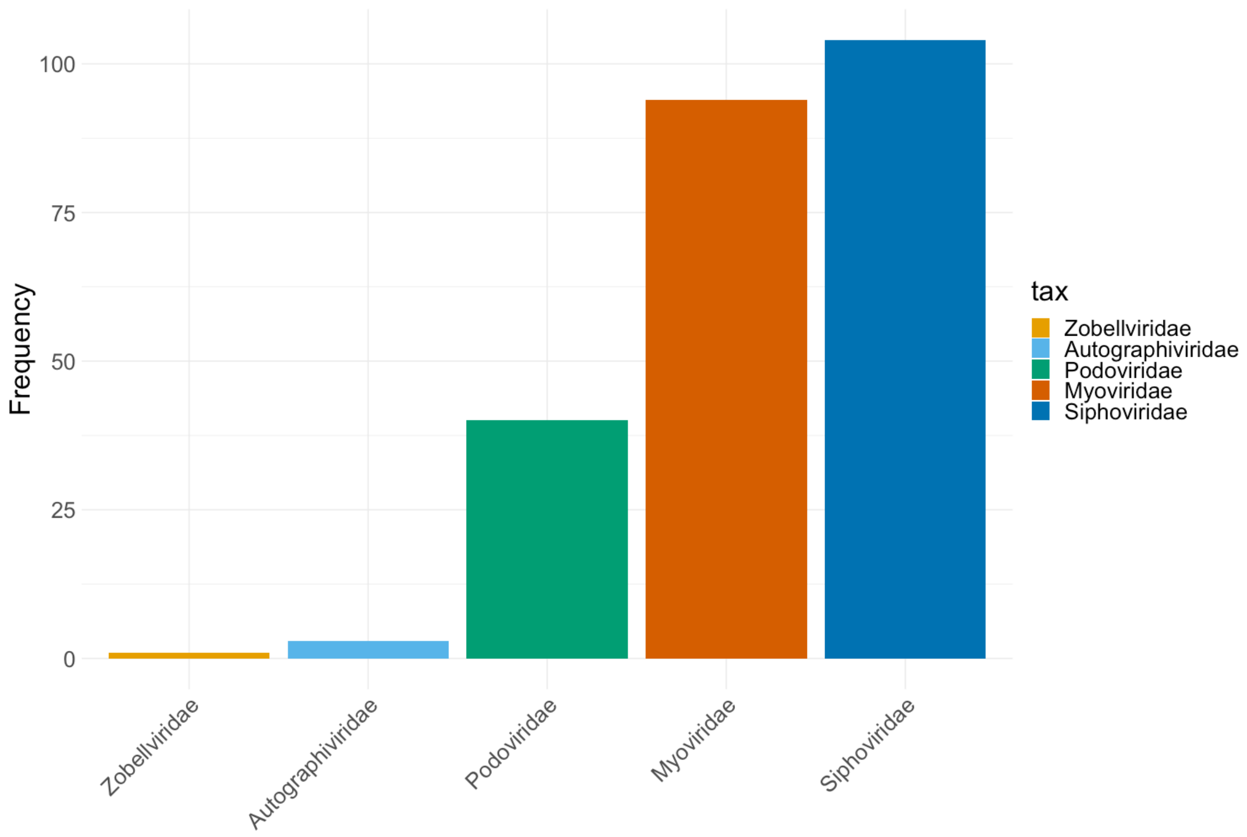


**Figure S4:** Bar plots showing the number of vOTUs successfully assigned to different phage families.

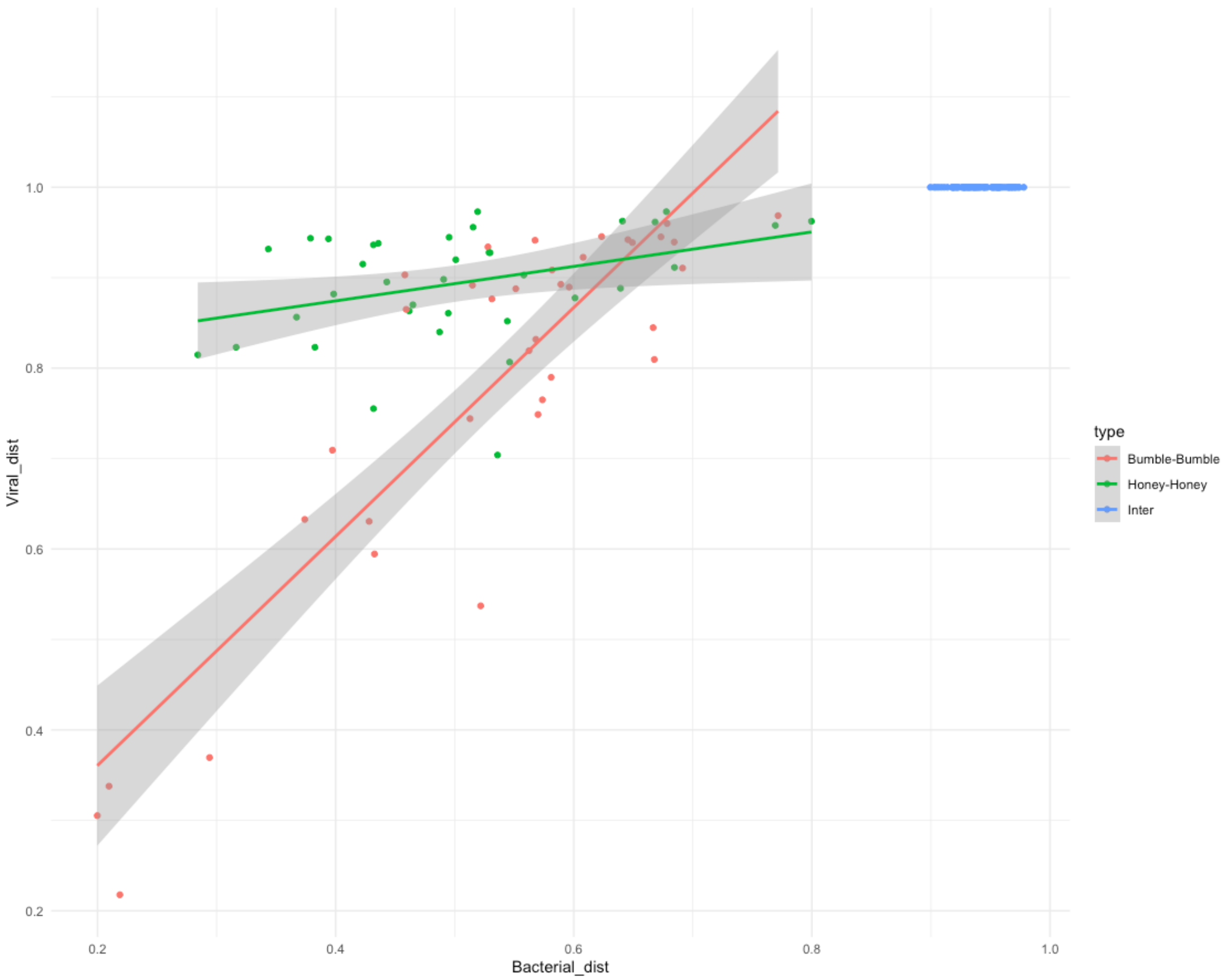


**Figure S5:** Scatterplot visualizing the mantel test performed. Each point represents the pairwise distance between two samples in beta diversity space. The x-axis measures bacterial community distance. The y-axis measures viral community distance. Points are colored based on whether the sample pairs being compared are both bumble bees (red), honey bees (green) or two different types of bees (blue). There is a positive relationship between phage and bacterial community structure.

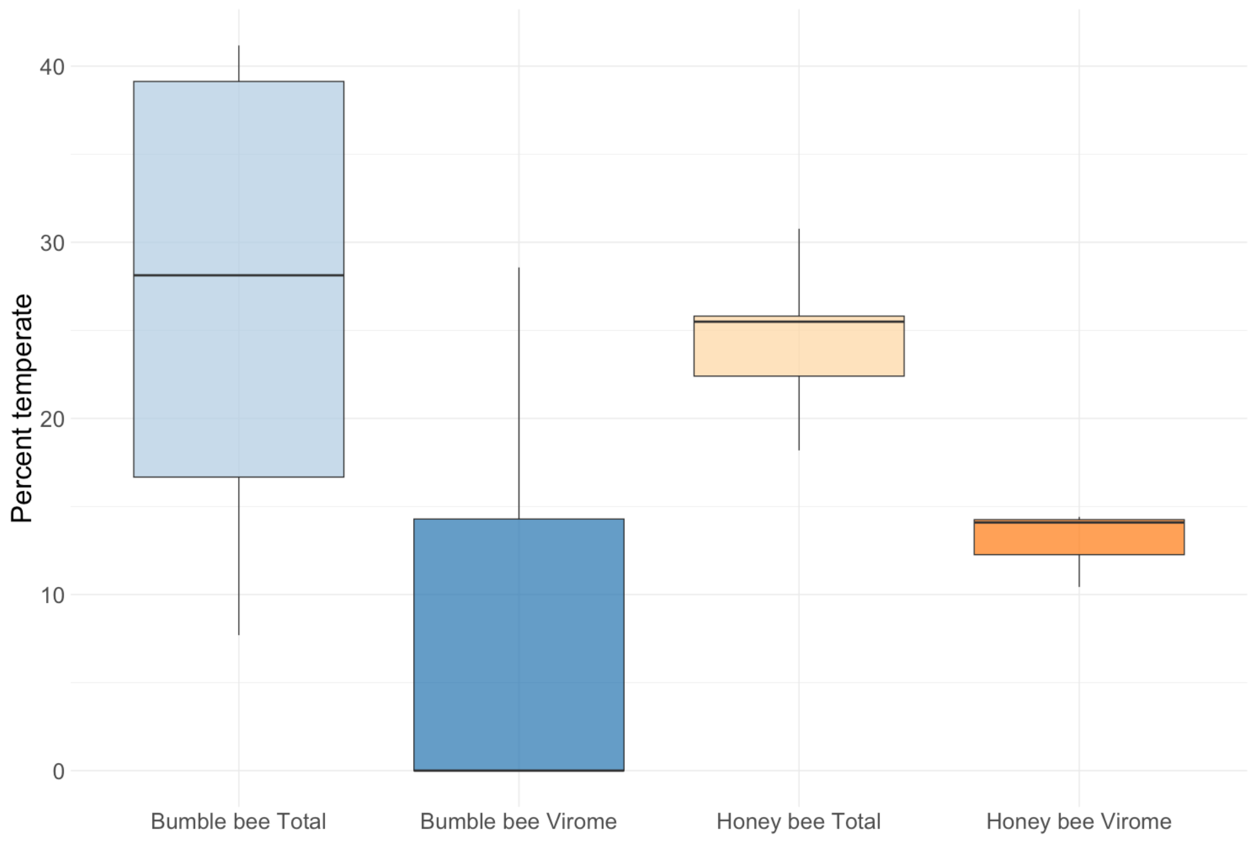


**Figure S6:** Boxplots describing the percent of vOTUs in each sample type predicted to be temperate. The x-axis describes sample group, the y-axis describes percent temperate vOTUs.

| | Bumble bee | | Honey bee | |
|---|---|---|---|---|
| | Total | Virome | Total | Virome |
| Acinetobacter | 0 ± 0[a] | 0 ± 0[a] | 0 ± 0[a] | 0.13 ± 0.09[b] |
| Lactobacillus | 30.07 ± 8.18[a] | 87.04 ± 7.17[b] | 21.07 ± 5.98[a] | 17.03 ± 1.84[a] |
| Gilliamella | 6.73 ± 1.25[a] | 0 ± 0[a] | 20.09 ± 4.25[a] | 8.81 ± 2.65[a] |
| Bifidobacterium | 4.84 ± 1.91[a] | 2.63 ± 1.53[a] | 4.01 ± 1.35[a] | 24.3 ± 2.34[b] |
| Bombilactobacillus | 5.78 ± 4.51[a] | 0 ± 0[a] | 0.16 ± 0.16[a] | 0.17 ± 0.01[a] |
| Schmidhempelia | 4.26 ± 0.43[a] | 0 ± 0[b] | 0 ± 0[b] | 0 ± 0[b] |
| Other | 0.11 ± 0.11 | 0 ± 0 | 0.03 ± 0.03 | 0.13 ± 0.06 |
| Unassigned | 47.53 ± 8.09 | 10.32 ± 6.44 | 54.43 ± 2.59 | 48.64 ± 5.67 |

**Table S1:** Table describing the distribution of phage predicted to target common bee associated bacteria. Letters represent results of emmeans posthoc test.
