## Supplemental Methods for "Targeted Viromes and Total Metagenomes Capture Distinct Components of Bee Gut Phage Communities"

### Bee rearing, sampling, and experimental design

The primary goals of this research were to 1) compare how targeted viromes and total metagenomes sample bee gut phage communities, and 2) compare the phage communities hosted by honey and bumble bees. To accomplish these goals, we sampled three colonies of managed honey bees and three colonies of commercially reared bumble bees.

Commercially reared *B. impatiens* were purchased from Koppert Biological Systems (Howell, MI, USA) and maintained in the laboratory at ambient conditions. Bumble bee colonies were provided with approximately 20g nonsterile honey bee collected pollen (Koppert Biological Systems) every 2-3 days and had access to artificial nectar (Koppert Biological Systems) *ad libitum*, as per manufacturer's recommendation. Honey bees (*A. mellifera*) were sampled from colonies maintained at the University of California, Davis and were collected between January 30$^{th}$ – February 13$^{th}$ 2023, 8:00am – 10:00am. Honey bees were sampled from the inside of colonies by removing the top of hive boxes and vacuuming bees with a handheld bee vacuum. From each colony, we generated one targeted virome and three total metagenomes (**Fig. 1**). To generate the large biomass required for targeted viromes, the guts of 100 bees (all sampled from the same hive) were pooled, as has been done previously[1,2]. Total metagenomes were generated from the guts of individual bees.

Though there are no official Institutional Animal Care and Use Committee (IACUC) guidelines for insects, we sought to euthanize our bees as humanly as possible. To this end, bees were first anesthetized via a 60 sec $CO_2$ exposure, followed by decapitation of individual bees. Immediately, the mid-hind gut section of 100 bees (all sampled from the same colony) were dissected and pooled for same day targeted virome extractions. Bees not dissected on the day of euthanasia were stored at -20°C pending mid-hindgut dissection and total metagenomic DNA extraction on the following day. All dissections took place in sterile PBS using ethanol and flame sterilized forceps. Each bee colony generated one targeted virome and three total metagenomes (**Fig. 1**).

Sample preparation and DNA extraction

Phage enrichments for targeted viromes were carried out following a protocol adapted from Goller et al[3], Santos-Medellin et al[4], and ter Horst et al[5] (https://www.protocols.io/view/soil-viromics-protocol-emerson-lab-v1-kxygxz7q4v8j/v1). Pools of bee gut material were combined with 10 ml of Protein-supplemented PBS (PPBS: 2% bovine serum albumin, 10% phosphate-buffered saline, 1% potassium citrate, and 150 mM MgSO4 in ultrapure water) and homogenized in a FastPrep-24 5G Homogenizer (MP Biomedicals, Irvine CA, USA) for 20 seconds at 6 m/s. Homogenates were then submitted to a series of washes to elute phage and other virus-like particles.

Washes began by centrifuging samples at 3,000 x g for 10min at 4 °C. The supernatant of this first wash was then collected in a new sterile 50 ml conical tube and stored at 4 °C. The remaining pellet of bee gut material was resuspended in 10 ml PPBS and recentrifuged at 3,000 x g for 10min at 4 °C. This second wash supernatant was then combined with the previously collected supernatant and stored at 4 °C. This was repeated one more time, for a total of three washes and 30 mL final supernatant volume.

Cells and small non-phage particles were then removed from samples through centrifugation and size filtration. The ~30 ml of phage elute produced by the previous washes was centrifuged at 10,000 x g for 15min at 4 °C and then filtered successively through an autoclaved glass fiber pre-filter and sterile 0.22 um PES vacuum filter.

Phage were pelleted by ultracentrifugation at 35,000 x g for 3 h at 4 °C and pellets incubated overnight at 4 °C with 200 uL of molecular grade water. Phage pellets were fully resuspended the following day via pipetting. Finally, to remove DNA not protected by a phage capsid, samples were combined with DNase (20 uL DNase, 20 uL DNase buffer) and incubated at 37 °C for 90 min. Reactions were terminated by adding of 20 uL of EDTA and incubating samples at 65 °C for 10 min.

To generate total metagenomes from individual bees, dissected guts were placed directly into bead beat tubes. DNA was extracted from both sample types using a DNeasy PowerSoil Pro kit (Qiagen, Hilden, Germany) following the manufacturer's instructions with the optional Qiagen Vortex Adapter 15 min bead beating step.

Read generation, pre-processing, and decontamination

Extracted DNA was submitted to the University of California Davis Genome Center for library prep and paired end 150 bp shotgun metagenomic sequencing. Library prep was performed using a Kapa Hyper prep kit (Kapa Biosystems-Roche, Basel, Switzerland) with Illumina TruSeq adapters (Illumina, San Diego, CA) and a target insert size of 350 bp. Sequencing took place on an Illumina NovaSeq. All samples were sequenced on the same run.

All sequencing data was trimmed and quality filtered the same way. Demultiplexed *.fastq.gz* files from the Davis Genome Center were processed using Trim-Galore[6] to remove polyG sequences. Next, we removed adapters, reads with an average quality <25, and reads smaller than 40 bp using Trimmomatic[7]. Contaminants and host sequences were then removed.

We first used metaSPADES[8] to construct contigs from negative sample reads. Negative sample contigs were then concatenated with the genomes of *A. melifera* (GCF_000002195.1) and *B. impatiens* (GCF_000188095.3). Lastly, Bowtie2[9] was used to build a database and filter high-quality reads.

<u>Computational phage identification</u>

To identify phage in both our targeted viromes and total metagenomes, metaSPADES was used to construct assemblies from processed reads. Contigs ≥5 kb were then selected using the BBDuk[10] toolset. Next, we used Virsorter2[11] to extract sequences ≥5 kb which had a strong (≥0.90) phage confidence. Lastly, BBDuk was used to rename sequences as Virome_phage_### and Total_phage_### based on samples they were recovered from.

<u>vOTU clustering, annotation, and table geneartion</u>

Viral OTUs (vOTUs) were constructed from putative phage sequences by combining all virome and total metagenome phage into a single *.fasta* file and clustering at 95% nucleic acid identity across ≥80% of the shorter aligned sequence using CD-HIT[12]. vOTUs were annotated with Prokka[13] using the PHROGs database[14]. The subsequent *.faa* files produced by Prokka were then parsed to assess vOTU gene content. vOTUs were classified as temperate when they were predicted to encode an integrase gene.

Finally, to estimate the abundance of phage within each sample, high-quality decontaminated reads were mapped against a local database containing all vOTU sequences using CoverM[15] trimmed_mean. Reads were considered mapped if ≥75% of the read mapped back to the phage sequences with a percent identity ≥95%. To only examine activate phage, coverage values <5 were considered 0. The resulting coverage table was then normalized to coverage per million reads.

Statistical and ecological analyses

Statistical and ecological analyses were conducted in R[16] v4.2.3. To determine how richness and diversity of phage communities differed in response to host bee, sampling method, and their interaction, we used lme4::lmer()[17] and lmerTest::lmer()[18] to construct linear mixed effect models. Phage richness was calculated by converting our vOTU table into a presence/absence table, and then taking the sum of phage in each sample. Shannon's diversity was calculated using vegan::diversity(index = "shannon")[19]. Lastly, emmeans::emmeans()[20] was used as a posthoc test.

To perform our rarefaction analysis, we used the seqtk[21] toolkit to randomly subsample cleaned virome read libraries. Up to 3.5 million read pairs were subsampled from each library in increasing increments of 100,000. The program coverm was then used to map subsampled reads back to vOTU sequences and calculate coverage values. Resulting coverage tables were processed following the same parameters indicated above (reads considered mapped if ≥75% of read mapped to phage sequence with ≥95% ANI, vOTU coverage values <5 were considered 0).

To explore how sampling method or bee species affected community structure, we used vegan::metaMDS(distance = "bray", k=2, trymax=1000) to calculate Bray-Curtis dissimilarities among samples. We visualized these differences using Non-metric Multidimensional Scaling ordination. We then tested if sampling method, bee species, or their interaction were associated with variation in community structure using the PERMANOVA test vegan::adonis(method="bray"). Colony ID was included as a random effect.

Bacterial community and density measurements

Bacterial communities were predicted directly from cleaned total metagenomic reads with Kraken2[22] and Bracken[23] using default parameters. Bacterial copy number was quantified via qPCR of the same total metagenome DNA extracts used in sequencing. Per reaction, master mix solutions contained: 5ul SSO Advanced Universal SYBR Supermix 41 (Catalog# 1725271), 0.3ul of both forward (799F= 5'-AACMGGATTAGATACCCKG-3') and reverse (1115R= 37 5'-AGGGTTGCGCTCGTTG-3') primers (10uM), 3.4ul Molecular grade water, and 1ul of template DNA (diluted 1:1000 in molecular grade water). Reactions were performed in triplicate for each sample.

Network generation

To compare the similarity of recovered phage from this and previous bee-phage studies[1,24,25], we used a gene-sharing network approach[26]. Gene-sharing networks were generated using the program vConTACT2[27,28]. The *gene2genome.tsv* file required by this program was constructed using custom scripts (supplementary S#). Briefly, vOTUs from our study and previously described bee phage sequences[1,24,25] were collected into a single directory and annotated with Prokka using PHROGs. Within each *.faa* file, genes were renamed to encode phage and gene number. Finally, all genes were concatenated into a single file and a for-loop run to parse protein ID, phage ID, and gene annotation as separate columns. Two phage from our study (Total_255 and Total_185) and four phage from Busby et al (Busby_72, Busby_91, Busby_123, Busby_126) failed to create *.faa* files in prokka (perhaps due to small genome size) and were excluded from our network analysis.

To produce gene-sharing networks, vConTACT2 was run with and without a reference database. The run with the reference database was performed to predict phage taxonomy, while the run without a reference database was conducted to produce the networks shown in this publication, as per Rosso[1] and Busby et al[24]. In both cases, vConTACT2 was run in "Diamond" rel-mode using the Markov clustering algorithm for protein clusters and the ClusterONE algorithm for viral clusters, as specified by Rosso et al[1].

Phage host prediction

To assigned putative bacterial hosts to detected phage, we performed a CRISPR-spacer analysis which pairs bacterial genera with phage which may infect them. For this, we first built bacterial metagenome assembled genomes (MAGs) from our total metagenomic dataset. The program Bowtie2 was used to map high-quality total metagenomic reads to the total metagenome assemblies produced by metaSPADES. Resulting *.sam* files were converted to *.bam* and sorted using Samtools[29]. Then, MetaBat2[30] and CheckM[31] were used to bin assemblies and quality check MAGs. If MAGs were predicted to have ≥2% contamination or be ≤90% complete, they were removed. Lastly, we used dRep[32] to remove redundant MAG bins.

To augment our host assignments, we then searched NCBI for assemblies of bacteria previously described in the bee gut. Collected assemblies were filtered to only retain those with: completeness ≥90%, contamination ≤2%, N50 ≥25kb, and coverage ≥30x. If any of these values were listed as "null" by NCBI, they were considered 0. Assemblies collected from NCBI were then combined with the MAGs generated in our study. CRISPR spacers were extracted from all assemblies using MINCED[33] and aligned to vOTUs using Blastn[34]. Only CRISPRs which mapped with 100% identity were considered aligned. Phage which recruited no CRISPRs were considered “unassigned”.

Lastly, to improve the number of vOTUs with host assignments, we combined the individual phage-host assignments generated by our CRISPR analysis with the clusters predicted by our vConTACT2 network to infer the hosts of whole clusters, as was done by Rosso et al[1]. If all vOTUs in a cluster were assigned the same host, that host was assigned to that cluster. If a cluster contained unassigned vOTUs and a single predicted host, then the single predicted host was applied to all vOTUs in the cluster. Lastly, if a cluster contain more than one assigned host, the vOTUs in that cluster retained their original host assignments.
